# BiCoN: Network-constrained biclustering of patients and omics data

**DOI:** 10.1101/2020.01.31.926345

**Authors:** Olga Lazareva, Hoan Van Do, Stefan Canzar, Kevin Yuan, Jan Baumbach, David B. Blumenthal, Paolo Tieri, Tim Kacprowski, Markus List

## Abstract

**Motivation:** Unsupervised learning approaches are frequently employed to identify patient subgroups and biomarkers such as disease-associated genes. Thus, clustering and biclustering are powerful techniques often used with expression data, but are usually not suitable to unravel molecular mechanisms along with patient subgroups. To alleviate this, we developed the network-constrained biclustering approach BiCoN (Biclustering Constrained by Networks) which (i) restricts biclusters to functionally related genes connected in molecular interaction networks and (ii) maximizes the difference in gene expression between two subgroups of patients.

**Results:** Our analyses of non-small cell lung and breast cancer gene expression data demonstrate that BiCoN clusters patients in agreement with known cancer subtypes while discovering gene subnetworks pointing to functional differences between these subtypes. Furthermore, we show that BiCoN is robust to noise and batch effects and can distinguish between high and low load of tumor-infiltrating leukocytes while identifying subnetworks related to immune cell function. In summary, BiCoN is a powerful new systems medicine tool to stratify patients while elucidating the responsible disease mechanism.

**Availability:** PyPI package: https://pypi.org/project/bicon

Web interface: https://exbio.wzw.tum.de/bicon

**Supplementary information:** Supplementary data are available at *Bioinformatics* online.

## 1 Introduction

Biomarkers are essential for stratifying patients for diagnosis, prognosis, or treatment selection. Currently, individual or composite molecular biomarkers based on, e.g., expression, methylation, mutation status, or copy number variation are used. Biomarker discovery has greatly benefited from supervised methods that identify molecular features that have a strong association with disease-relevant variables such as drug response, relapse, survival time, or disease subtype. However, supervised methods are strongly biased by our current understanding of diseases, in particular by disease definitions that were established before rich molecular data became available. While classical unsupervised methods such as clustering have been successfully applied in the past, e.g., to reveal gene signatures predicting breast cancer subtypes (Parker *et al.*, 2009; Nielsen *et al.*, 2010), they group patients based on the entire molecular profile, and overlook meaningful differences limited to a subset of genes.

Biclustering aims to discover rows in a matrix which exhibit similar behaviour across a subset of columns and *vice versa* (Hartigan, 1972). It is suited for identifying disease-associated genes from gene expression data while stratifying patients at the same time (Prelic, 2006). As an NP-hard problem (Tanay *et al.*, 2002), biclustering is typically solved via heuristics. A gene expression matrix describes the expression of genes (rows) across samples (columns), which can reflect individual patients, time points or conditions. In patient stratification (i.e. splitting patients into clinically relevant subgroups), samples typically stem from different patients with a disease phenotype. In gene expression data, a bicluster defines a set of genes and a set of patients for which these genes are co-expressed (Cheng and Church, 2000). Gene co-expression does not imply a direct functional connection and, hence, genes identified by biclustering are often difficult to interpret. In contrast, molecular interaction networks such as protein-protein-interaction (PPI) networks capture direct and functional interactions. To obtain functionally coherent features sets, we extract sets of genes that form biclusters according to the expression data and are also connected in a molecular interaction network.

Many diseases are caused by aberrations in molecular pathways or modules of functionally related genes (Berg *et al.*, 2002). This suggests to focus on gene modules for delivering more interpretable and robust mechanistic explanations of disease phenotypes. Network enrichment methods leverage prior information of molecular interactions for identifying gene modules as subnetworks (Batra *et al.*, 2017). Gene modules are robust features for classification and disease subtyping (Alcaraz *et al.*, 2017). Few methods exist that can utilize molecular interaction networks for patient stratification. Two integer linear programming methods were suggested (Yu *et al.*, 2017, Liu *et al.*, 2014) both of which rely on the GeneRank (Morrison *et al.*, 2005) algorithm to incorporate network information. GeneRank depends on a parameter *θ* describing the influence of the network whose choice is not straight-forward and was shown to have a notable impact on the results (Yu *et al.*, 2017). None of the above methods actively encourage connected subnetworks as solutions and are thus not suited for discovering disease modules with mechanistic interpretation. To overcome this issue, we developed BiCoN, a network-constrained biclustering approach which delivers meaningful results on real-world datasets on par with other state-of-the art methods. We have validated our results on breast cancer (TCGA Pan-Cancer) and non-small cell lung carcinoma (NSCLC) datasets (Rousseaux *et al.*, 2013) and found that BiCoN is robust to batch effects and delivers biologically interpretable mechanistic insights into disease subtypes.

## 2 Approach

### 2.1 Problem statement

We expect that if gene expression shows opposite behaviour in the two subgroups (i.e. similar to conventional differential expression analysis), a biological function jointly carried out by these genes is active in one patient group and inactive in the other one. This assumption is reflected in our objective function and formally described below.

Consider a matrix of expression values *X*^*n*×*m*^ with *n* genes and *m* patients as well as *G* = (*V, E*), a molecular interaction network of gene set *V*. We further consider *P* as the set of *m* patients (samples) and construct a bipartite graph *B* with genes *V* and patients *P* as node types connected by weighted edges (*v, p*). Edge weights reflect the expression strength for a given patient from expression values *X*^*n*×*m*^. We construct a joint graph *J* by mapping *G* onto *B* via the shared genes in *V*. Our goal is to partition *P* into clusters *P*_1_, *P*_2_, and to find 2 connected subnetworks *G*_1_(*V*_1_, *E,*_1_), *G*_2_(*V*_2_, *E*_2_) of minimal size *L*_*min*_ and of maximal size *L*_*max*_. Size constraints can be adapted by users to the expected size of the molecular pathways, i.e. small subnetworks will represent more specific and large subnetworks more general molecular functions or biological processes.

Thus, we aim to derive patient groups (clusters) which are characterised by maximally differential expression in the extracted subnetworks:

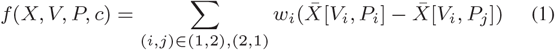

Where 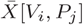 is the average expression of genes of module *i* for patients in cluster *j*, *w*_*i*_ is a weight for *G*_*i*_(*V*_*i*_, *E*_*i*_) which penalizes too small or too large, disconnected solutions:

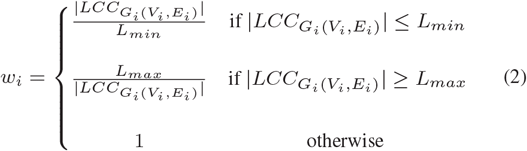

Where 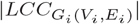 is the size of the largest connected component (LCC) in a subnetwork *G*_*i*_(*V*_*i*_, *E*_*i*_). Thus, *w*_*i*_ is always equal to 1 if the size of LCC corresponds to the user defined *L*_*min*_ and *L*_*max*_ and *w*_*i*_ < 0 means that the obtained solution does not fit into the desired range. Smaller *w_i_* means larger deviation from a user’s preferences.

To obtain more than two clusters, BiCoN can in principle be applied recursively to further split clusters as also shown in the application in Section 4.2.

### 2.2 BiCoN algorithm

BiCoN is a heuristic algorithm that aims to find differentially expressed subnetworks that can mechanistically explain patient stratification. This combinatorial problem can be addressed by various metaheuristic frameworks such as e.g. Genetic Algorithm (Banzhaf *et al.*, 1998) or Swarm Intelligence (Eberhart and Kennedy, 1995). We have chosen Ant Colony Optimization (ACO) (Stützle, 2009) as the main framework that performs exploration of the search space and Local Search (Aarts *et al.*, 2003) to ensure local optimality of the final solution. The combination of ACO and Local Search was shown to be very efficient in finding near-optimal solutions to hard combinatorial optimization problems (Stützle and Hoos, 1999) and leads to significant improvements compared to ACO or local search alone (Stutzle and Hoos, 1997). As we already had good prior experiences with ACO on similar problems (Alcaraz *et al.*, 2012) we expected that combination with local search will lead to high quality results.

ACO is a nature-inspired probabilistic technique for solving computational problems which can be reduced to finding optimal paths through graphs. We use ACO to identify a set of relevant genes for each patient which we subsequently aggregate into a global solution. A full description of the algorithm can be found in the Supplementary Material, section “Algorithm description” and the pseudo-code can be found below (Algorithm 1). We also describe the full workflow on Figure 1. Briefly, ants travel the joint graph *J* in three phases which are repeated until convergence:

1. An ant performs a random walk within nodes that are highly connected to a patient-node and makes greedy choices according to the objective function (Equation 1) by choosing genes which are most relevant to a patient (orange edges on Figure 1 step 2). The probability of selecting a gene for a certain patient depends on the combined information from gene expression values (which are encoded in the heuristic information matrix) and the ant’s “memories” on whether the choice of this gene has led to a quality solution in the previous rounds (pheromone matrix). More details about the implementation can be found in Supplementary Material, section “Algorithm description”.
2. Afterwards, the selected genes are used for clustering patients and for extracting sub-networks relevant to each patient cluster (Figure 1 step 3 and 4). A candidate solution is evaluated by the objective function score.
3. The best solution is used for updating the pheromone and probability matrices for the next iteration.

**Fig. 1:**
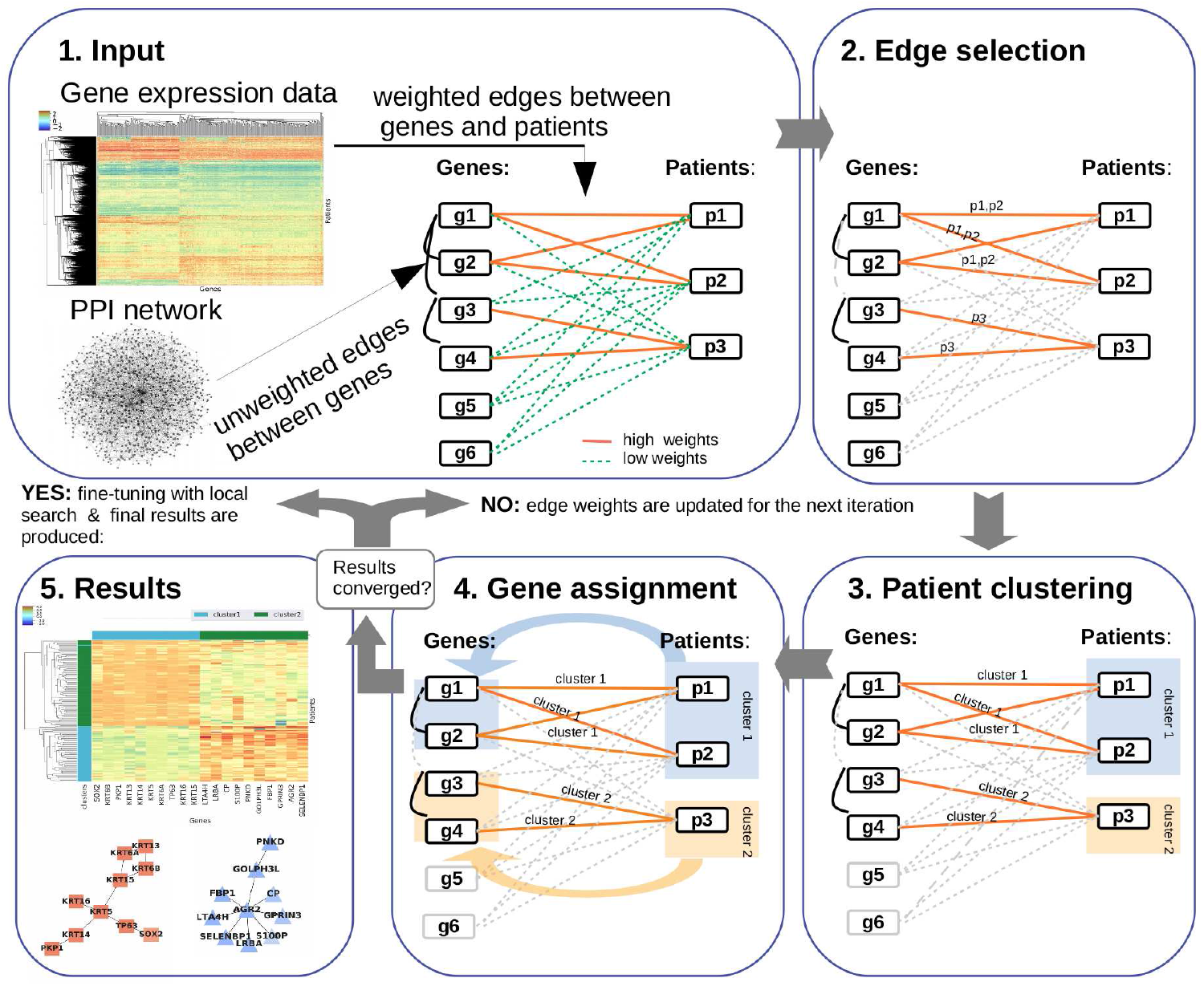
The algorithmic framework of BiCoN. (1) Gene expression data is converted to a bipartite graph and PPI interactions are added as edges between genes. (2) ACO is employed for feature selection (relevant edges) and subsequently patients (3) and genes (4) are clustered. Multiple possible solutions are computed in parallel and then evaluated and reinforced. As a result (5), BiCoN stratifies patients based only on subnetworks representing disease mechanisms.

When the best solution is obtained we perform local search for possible local improvements, i.e. iteratively apply changes to subnetworks (such as node insertion, deletion or substitution) and keep changes that lead to objective function maximization. This allows us to retrieve robust and stable solutions as well as to ensure local optimality.

Even though BiCoN uses several hyperparameters, our experiments have shown that those do not have a large impact on the results and the optimal combination is determined automatically based on the dimension and distribution of the expression matrix. Therefore, the user only has to specify the desired size of the solution subnetworks (*L*_*min*_ and *L*_*max*_).

## 3 Methods

### 3.1 Data collection and processing

#### 3.1.1 Gene expression data

TCGA breast cancer data was obtained through the UCSC Xena browser (https://xenabrowser.net/). The NSCLC dataset (accession number GSE30219, (Rousseaux *et al.*, 2013) was obtained using GEO2R (https://www.ncbi.nlm.nih.gov/geo/geo2r/). Both datasets were retrieved together with the corresponding metadata which contained annotated cancer subtypes.

For the NSCLC dataset, gene probes were mapped to Entrez gene IDs. If multiple probes corresponded to a single gene, the median value was used. We applied a *log*_2_ transformation to account for skewness of the data. Data was z-score transformed to indicate the magnitude of changes in gene expression in individual samples and conditions compared to the background. In most gene expression datasets, a majority of genes is lowly expressed and does not vary to a larger extend. To account for this and to improve run-time, BiCoN filters out genes with a small variance preserving only the *n* most variant genes (here *n* = 3000).

#### 3.1.2 Molecular interaction network

We used physical and genetic protein-protein interactions (PPI) in H. Sapiens from BioGRID (version 3.5.176). The network consisted of 343,563 unique interactions between 16,830 genes.

### 3.2 Simulation of Batch Effects

Batch effects are technical variations that have been introduced by external factors during handling of the samples (e.g. personnel effects, environmental conditions, different experiment times) (Luo *et al.*, 2010). While some of those effects can be minimised, batch effects are still almost inevitable in practice (Chen *et al.*, 2011). Thus, batch effect-resistance is very important for any modern algorithmic framework.

To demonstrate that BiCoN is robust to batch effects, we simulate data using a linear mixed effect model. We consider two variables: *cluster* and *batch*. The variable *cluster* indicates whether a gene is part of the foreground (*cluster* = 1 or *cluster* = 2) or the background (*cluster* = 0), i.e. it is not differentially expressed. The variable *batch* indicates the study or batch of expression values (*batch* = 1 or *batch* = 2). The expression values are simulated as follows:

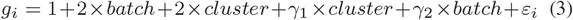

where the first part of the equation (1 + 2 × *batch* + 2 × *cluster*) is fixed and shared by all genes. Errors *ɛ*_*i*_ are independent and identically distribu ted (with zero mean). The random effects parameters *γ*_1_ and *γ*_2_ follow a bivariate normal distribution with zero mean, and variance 1 and 2 respectively, i.e. the technical variance is twice the biological variance.

**Algorithm 1:**
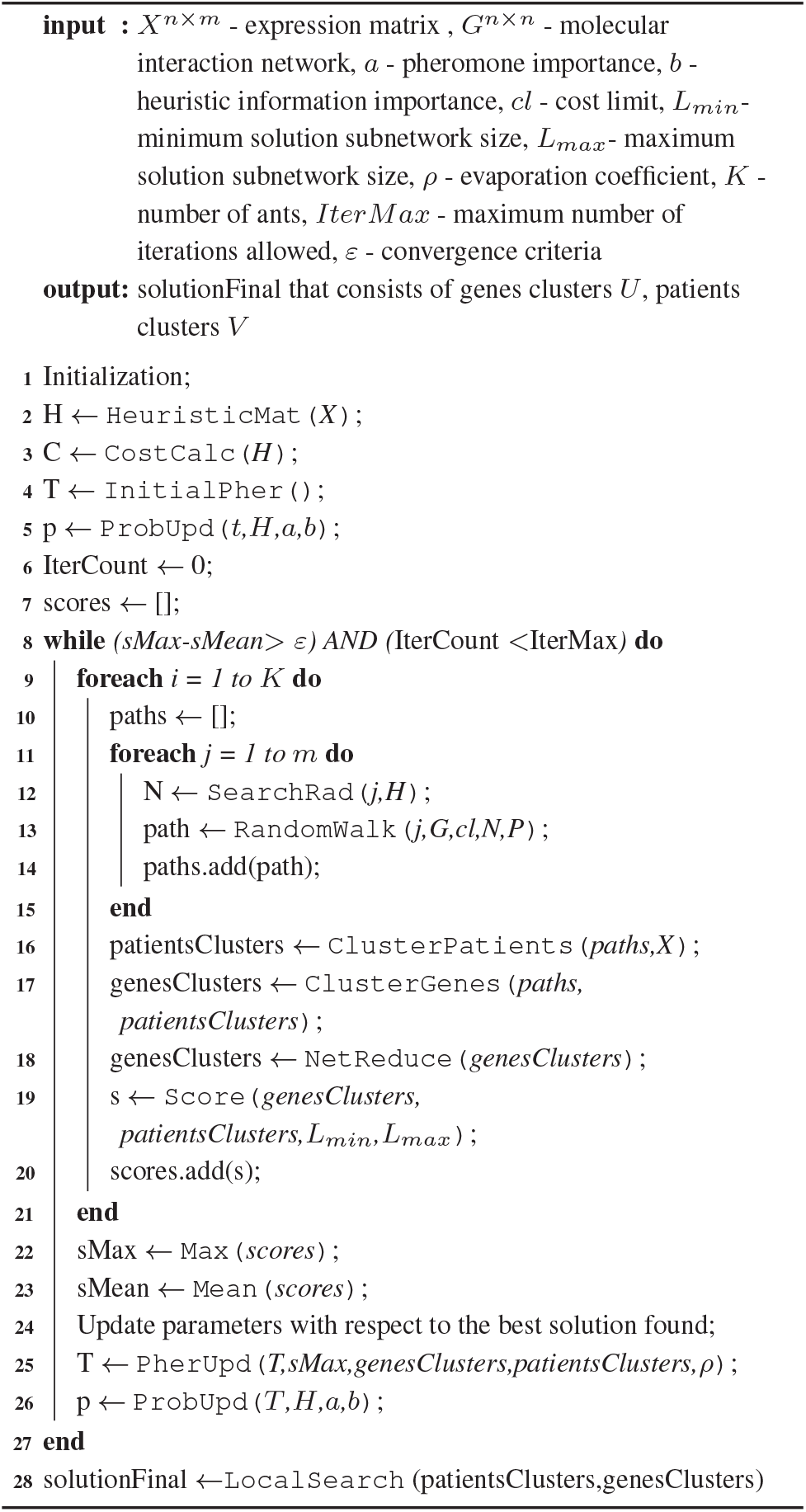
BiCoN

### 3.3 Benchmarking

To show how BiCoN results compare to commonly used clustering and biclustering algorithms, we performed multiple assessments for each of the methods listed in Table 1:

- To show how BiCoN can recover PAM50 annotated breast cancer subtypes (using TCGA data as a source), we computed Jaccard index (an intersection of two sets over the union) between the known subtypes and the resulting patients clusters/biclusters.
- To show how BiCoN can handle batch effect in comparison to other methods, we simulated data as described in section 3.2 and computed the overlap between known classes of patients and the resulting clusters/biclusters. To avoid favouring the assumption of genes connectivity used by BiCoN, we also repeated the simulation such that the signal-carrying foreground genes are randomly distributed over the network.

**Table 1.** Used algorithms

| Algorithm | Implementation | Parameters | Data processing applied |
| --- | --- | --- | --- |
| Bimax (Prelić *et al.*, 2006) | R package ‘biclust’ | minc:100, number:4 | normalization, binarization |
| QUBIC (Li *et al.*, 2009) | R package ‘biclust’ | o:4 | normalization, qualitative discretization |
| Plaid (Lazzeroni and Owen, 2002) | R package ‘biclust’ | row.release:0.5, col.release:0.9 | normalization, binarization |
| Spectral (Shi and Malik, 2000) | Python library ‘sklearn’ | n_clusters:2, affinity: “nearest_neighbors” | normalization |
| Hierarchical (Voorhees, 1986) | Python library ‘sklearn’ | n_clusters:2 | normalization |

Nearest neighbors affinity was selected as it produced results with the highest Jaccard index for both instances

As a metric for comparison we used Jaccard index rather than MCC as it allows to measure relationship between resulting biclusters and the actual classes even when the patients biclusters overlap and do not include all patients.

Even though we use classical clustering methods for benchmarking, we emphasise key differences between the suggested approach and classical clustering. BiCoN extracts biological mechanisms that explain patient stratification. Even though subnetworks extraction after clustering of patients is feasible, to our knowledge there is no gold standard for this procedure. While it is possible to extract subneworks and disease mechanisms subsequent to clustering or by relying on known disease subtypes (Alcaraz *et al.*, 2017), we argue that such clusters are driven by global differences and not by the activity of a single disease mechanism. Hence, extracting disease mechanisms along with patient stratification is better suited to identify patient subgroups affected by key disease mechanisms. Moreover, clustering performed on the whole genome is also not advisable as the use of multidimensional data can lead to multiple negative effects, which are often referred to as “curse of dimensionality” (Thangavelu *et al.*, 2019).

For all algorithms in Table 1, we chose parameters that maximize performance for each of the methods.

## 4 Results and Discussion

We evaluated BiCoN on simulated and real data with respect to the robustness of patient clustering and gene selection as well as robustness to batch effects. Furthermore, two application cases illustrate the practical use of BiCoN.

### 4.1 Robustness analysis

#### 4.1.1 Noise robustness

To introduce varying levels of noise to a data set, we randomly select between 0 and 90% of the genes and randomly permute their expression values. A noise level of 0.1 means that the expression vectors of 10% of genes were permuted. For each noise level, we average results over 10 independent runs.

We use the NSCLC data set with two annotated subtypes as gold standard: adenocarcinoma and squamous cell carcinoma. As evaluation metrics, we consider the value of BiCoN objective function as well as Matthews Correlation Coefficient (MCC) (Matthews, 1975) between the proposed clusters and cancer subtype labels. The latter is meant to demonstrate that BiCoN is able to recover cancer subtypes while inferring a mechanistic explanation for the subtype differences. For this analysis, we retain the 3000 most variant genes and set parameters *L*_*min*_ = 10 and *L*_*max*_ = 25 to control the size of the solution.

Figure 2(a) shows a consistent decline in the objective function with increasing noise, indicating that the algorithm is reacting reasonably to the decline in data quality. Figure 2(b) shows that the algorithm is able to recapture the cancer subtypes almost perfectly (average MCC higher than 0.9) up to a noise level of 0.5 where 50% of the data have been permuted. Figure 2(c) shows a strong positive correlation between the objective function value and MCC, which confirms that the objective function is high when cancer subtypes are well separated.

**Fig. 2:**
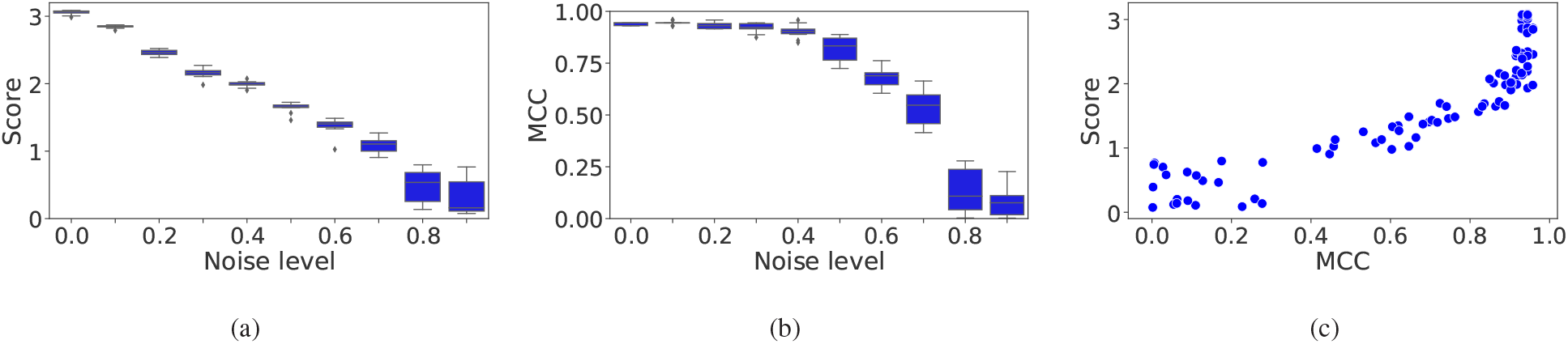
Robustness analysis. a) Objective function score versus the percentage of noisy data. b) Matthews Correlation Coefficient (MCC) with respect to the known classes versus the percentage of noisy data. c) Correlation of objective function scores and MCC.

#### 4.1.2 Batch effect robustness

Batch effects are a common problem in data analysis (Goh *et al.*, 2017) and many methods have been proposed for removing batch effects from data (Lazar *et al.*, 2012). However, removing batch effects may also remove biologically relevant group differences from the data. Batch effect correction methods that are designed to retain group differences can lead to exaggerated confidence in downstream analyses (Nygaard *et al.*, 2016). In unsupervised analysis, this issue is critical, since we, by definition, do not know the relevant sample or patient groups *a priori*.

BiCoN is a graph-based method and, hence, it is not as strongly affected by the global distribution of expression values as classical clustering methods. Pre-processing methods that scale data to a certain range enforce it to have certain mean and variance (e.g. z-scores) or make the distribution more symmetrical (e.g. *log*_2_ transformation) are not ideal for batch effect correction as they do not differentiate between signal and noise. In this scenario, a graph-based method benefits from the assumption that the joint signal of the genes in a subnetwork is stronger than that of individual genes.

To study if BiCoN can indeed tolerate batch effects, we simulate gene expression data (see Methods for details) where we introduce a batch effect with a larger variance than for the group difference. Our aim is to show that BiCoN can leverage the network to recover the signal even if it is overshadowed by batch effects.

We have simulated expression data for 2 × 20 foreground genes (two biclusters) and for 1000 background genes. We also tested the performance with 2 × 30, 2 × 40 and 2 × 60 foreground genes.

The network was simulated as three disjoint Barabasi-Albert graphs (one for each of genes biclusters and one for background genes) (Barabási and Albert, 1999) which were connected by random edges until they have reached the same density as the BioGRID network (0.0013). Barabasi-Albert graphs have similar node degree distribution as the BioGRID network and were thus considered suitable for the simulation study.

Figure 3(a) shows that the batches differ in their distribution, causing hierarchical clustering to group samples by batch rather than by disease phenotype. Figure 3(b) shows that differences due to batch effects are eliminated after z-score normalization. We can also see that the difference between the sample groups is now lost and can not be recovered by hierarchical clustering. Figure 3(c) shows that in spite of this noise, BiCoN can recover the disease phenotype together with the foreground genes. Thus, when two datasets can be normalized separately (e.g. z-scores are applied to each dataset), BiCoN is uniquely suited to cluster patients where individual gene modules are disturbed. Even when the signal is obscured by batch effects, the functional connection of solution genes in the network (Figure 3 (d)) helps to robustly recover the signal.

**Fig. 3:**
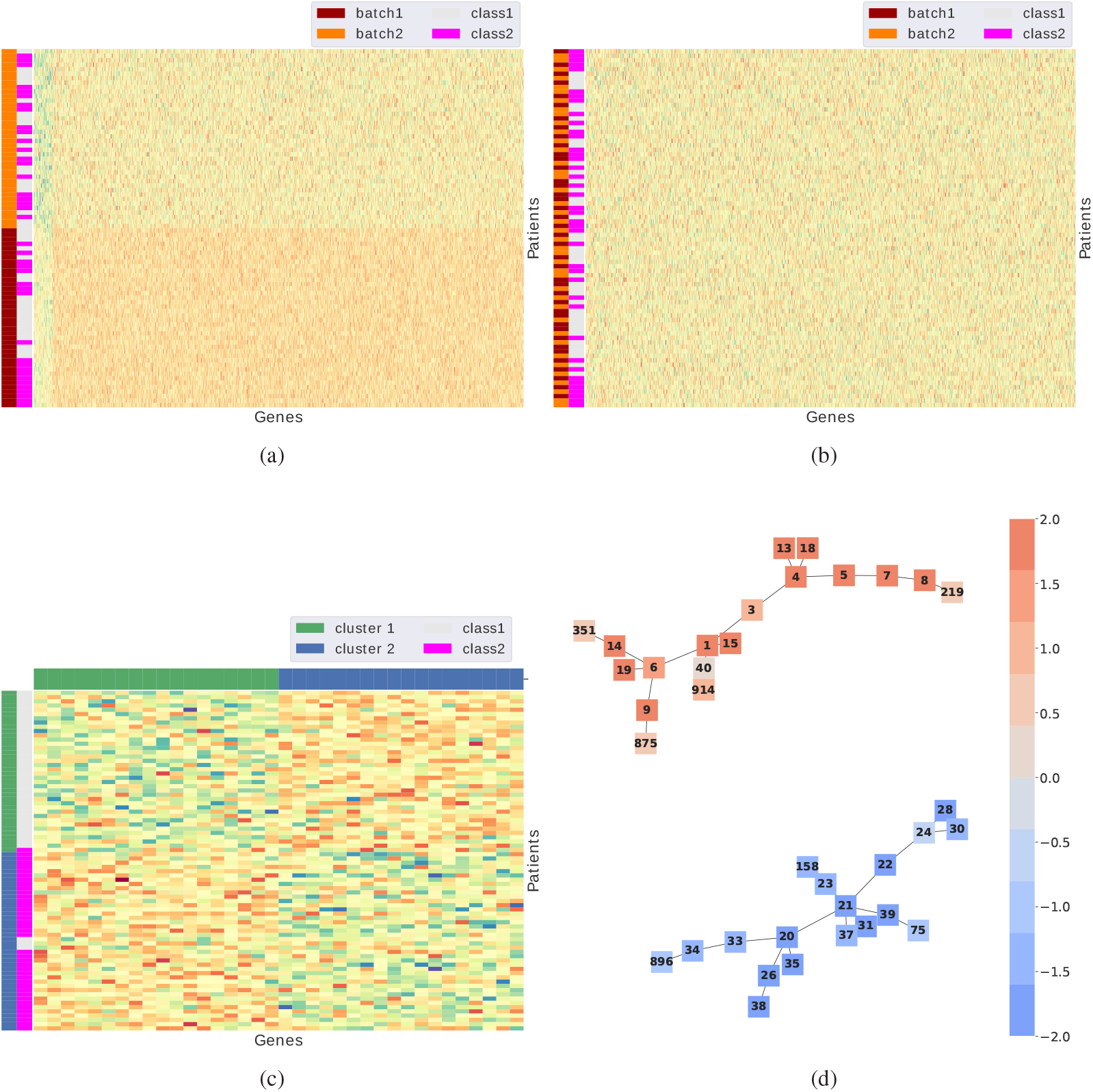
a) Two data sets with different distributions due to batch effects.b) The merged data sets after z-scores normalization. The batch effect vanishes, but the disease phenotype is still not distinguishable. c) BiCoN is able to recover the initial disease phenotypes with Jaccard index of 0.92 (in average after 10 runs) while extracting the 40 foreground genes out of 1000 background genes. d) The resulting subnetworks with all seeded solution genes are colored with respect to their average expression in one cluster.

To show how BiCoN results align with other clustering and biclustering methods, we have simulated 10 datasets with batch effect and evaluated the performance. To make sure that we do not put BiCoN in favour by enforcing connectivity of genes, we also performed simulations with a single Barabasi-Albert graph, where foreground genes were randomly distributed (Figure 4).

**Fig. 4:**
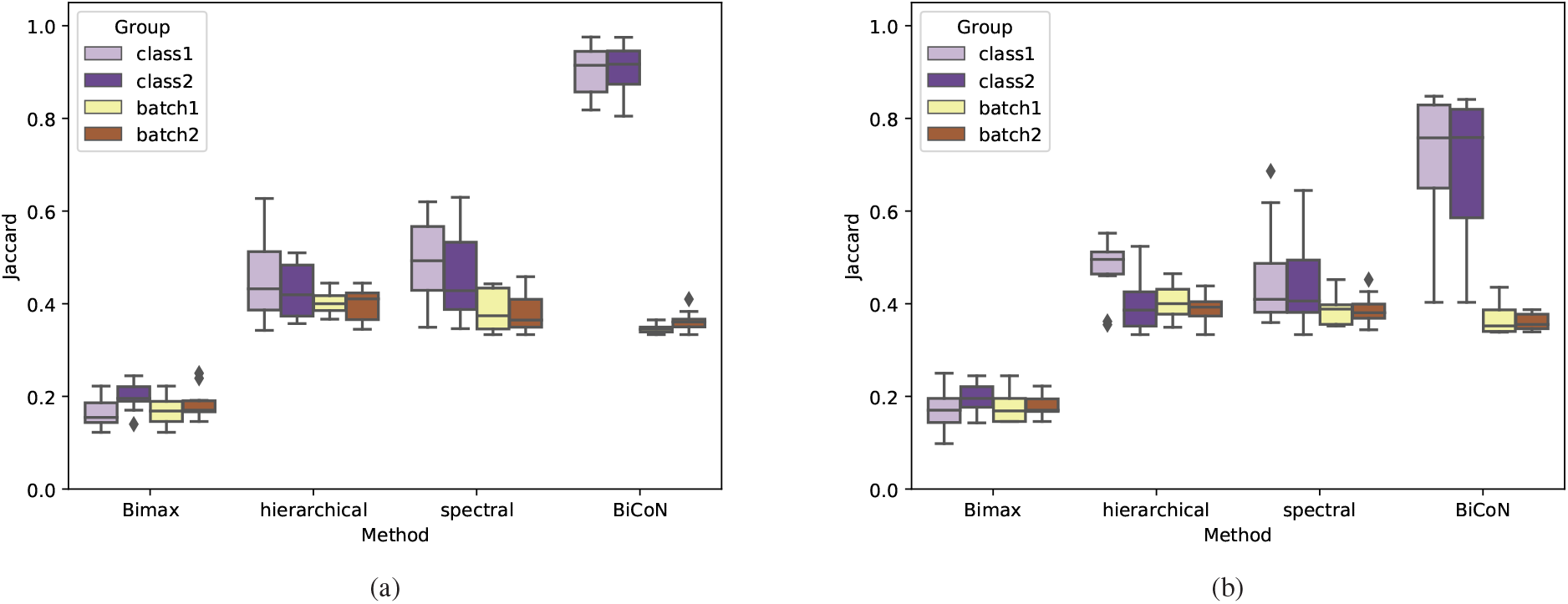
Jaccard indices between the patients clusters and actual subgroups (class 1 or class 2) as well as with batches of patients (batch 1 and batch 2) for 10 simulated datasets with a strong batch effect. a) When foreground genes are connected in a network, BiCoN clusters patients almost perfectly based on the actual signal. b) When the foreground genes are randomly distributed in the network, BiCoN still achieves higher performance than other methods that were capable to find any clusters. Plaid and QUBIC were not able to find any clusters and were excluded from further assessment.

Among the considered biclustering algorithms (Table 1), only Bimax was capable of finding any clusters, while Plaid and QUBIC could not find any structure in the given data regardless of chosen parameters and therefore was excluded from further assessment. The experiments showed that even though the quality of the results drops when the foreground genes are not directly connected, BiCoN still performs significantly better than other methods. This is due to the fact, that like a real PPI, the simulated network had power-law node degree distribution which means that the network diameter is rather small and therefore most of the nodes are still reachable through hub-nodes even when they are not directly connected. Thus, BiCoN performance dropped when using random networks but still outperformed other methods that are not network-restricted.

### 4.2 Application to TCGA breast cancer data

We applied BiCoN to the TCGA breast cancer dataset. We expected BiCoN to be able to recover known subtypes assigned via the PAM50 gene panel (Parker *et al.*, 2009; Nielsen *et al.*, 2010) which is a gold standard in breast cancer subtype prediction. For the analysis, we focused on patients with the most common molecular subtypes, luminal (estrogen-receptor and/or progesterone-receptor positive) and basal (hormone-receptor-negative and HER2 negative).

As a proof of concept, we first showed thatBiCoNcan separate patients into the two clinically well distinguishable subtypes luminal and basal. Next, we applied BiCoN separately for patients with luminal and basal subtype to investigate how patients are stratified in a more challenging scenario. For each subgroup, we ran the algorithm 10 times and selected a solution with the highest score based on the previous observation that the highest objective function score corresponds to the highest correlation between the resulting biclusters and the expected patient groups.

#### 4.2.1 Luminal versus basal separation

As expected, the separation between patients with luminal and basal breast cancer subtypes is straightforward. The clusters correspond to the subtype labels. In Figure 6(a), the separation between patients groups matches the PAM50 classification (average Jaccard index is equal to 0.99). BiCoN not only performs as well as methods like hierarchical clustering (Figure 5) (Jaccard index is equal to 0.96) but also yields two differentially expressed subnetworkswhich explain subtype differenceswith a vastly lower number of genes than a classical clustering method while offering a mechanistic explanation of subtype differences.

**Fig. 5:**
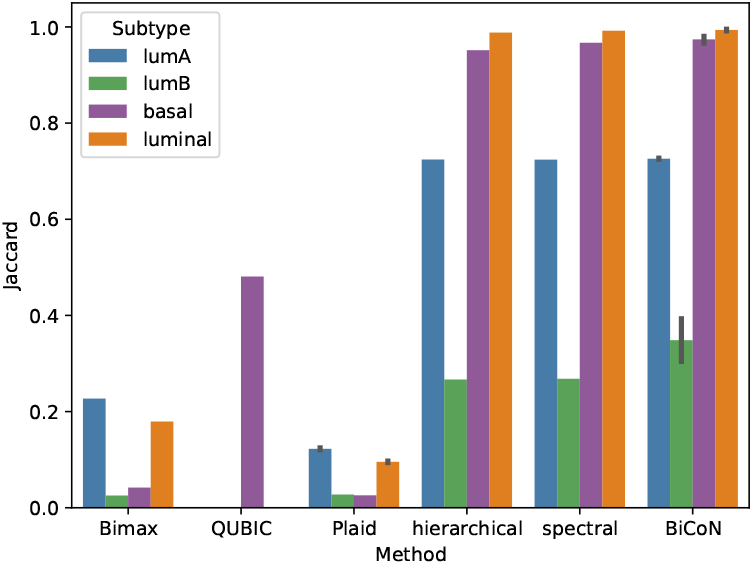
TCGA breast cancer subtypes identification by various algorithms (for 10 runs). Jaccard index was computed as a best match between produced patients clusters and the known breast cancer subtypes for BiCoN and other well-known clustering and biclustering algorithms. BiCoN shows performance which is comparable with other clustering algorithms while also revealing functionally connected subnetworks which explain the phenotype.

**Fig. 6:**
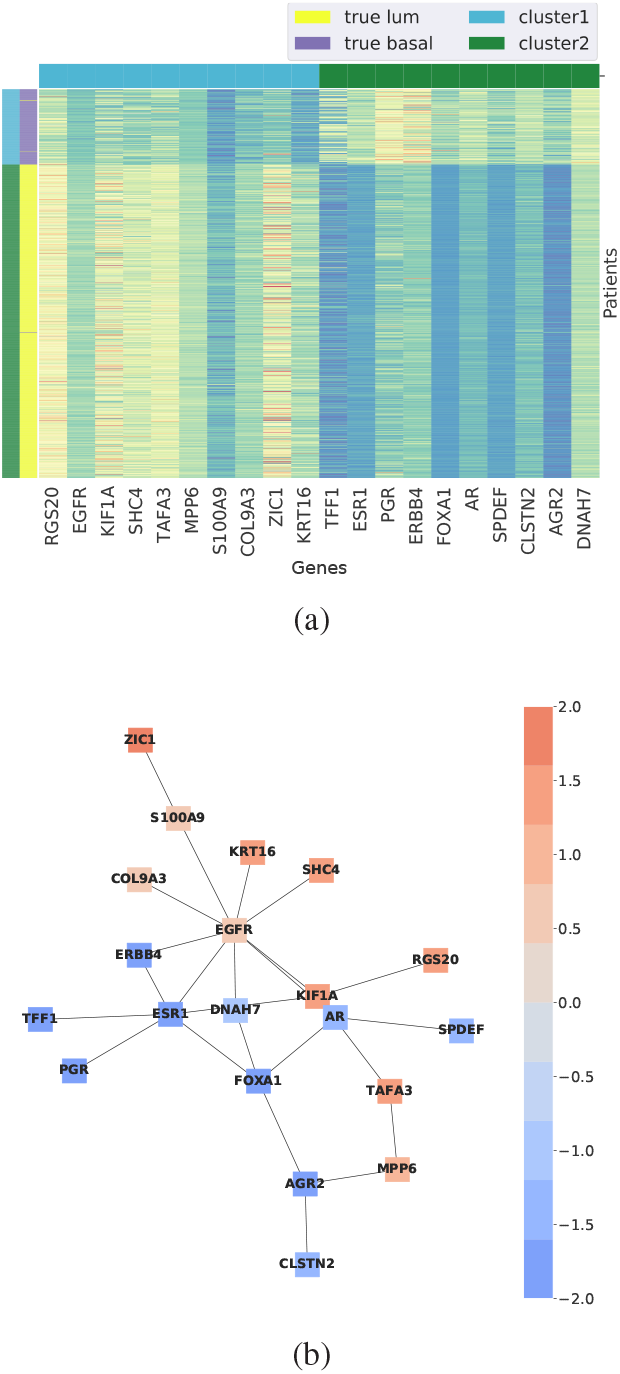
a) Biclusters for patients with luminal and basal breast cancer subtype. b) Resulting subnetwork: nodes are colored with respect to their mean expression in one of the clusters and thus nodes colored orange have very high average expression in the cluster and the blue nodes have very low expression values.

The extracted subnetworks in Figure 6(b) show a strong difference in expression. Note that while BiCoN restricts genes inside a bicluster to be connected, it does not impose any relationships between two biclusters. As a consequence, it is possible that the resulting subnetworks overlap.

In contrast to methods yielding gene signatures such as PAM50, BiCoN focuses on revealing specific pathways. Enrichment analysis of cancerrelated pathways (Supplementary Material Figure S3) confirms strong association of the resulting genes with breast cancer subtype-specific signalling, in particular estrogen signaling pathway (adjusted p-value = 0.018) and ErbB signaling pathway (adjusted p-value = 0.025).

Random-walks on scale-free networks are biased towards hub nodes since these have a high degree (Gillis *et al.*, 2014). BiCoN avoids this hub bias as it performs random walks on the joint graph of a PPI and expression data which is not scale-free. Consequently, the selected nodes have approximately the same degree distribution as the input network (Supplementary Materials Figure S1).

#### 4.2.2 Luminal patient stratification

Next, we consider only patients that were originally classified as luminal subtype to see if we can further stratify them into subtypes luminal A and luminal B which are known to be difficult to separate on the level of gene expression. Here, our solution does not completely agree with the PAM50 classes (mean Jaccard index 0.49), although we observe two clearly separable groups and that most of the luminal B patients were placed in cluster 1 (Figure 7). To investigate possible differences in cell type composition of the two clusters, we used the signature-based deconvolution method xCell (Aran *et al.*, 2017). xCell estimates contributions of 64 immune and stromal cell types. In addition, it provides aggregated scores such as an immune score, a stromal score and a microenvironment score. xCell is based on harmonized data from 1,822 purified cell type transcriptomes from various sources and employs a curve fitting approach for linear comparison of cell types. Clusters reported by BiCoN show significant differences between cell types scores. The strongest difference between patients is found in the stromal score (−*log*_10_ p-value is over*>* 55), hematopoietic stem cells (−*log*_10_ p-value > 50) and CLP cells (−*log*_10_ p-value > 50). See Supplementary Figures S4(a), S5(a) for details.

**Fig. 7:**
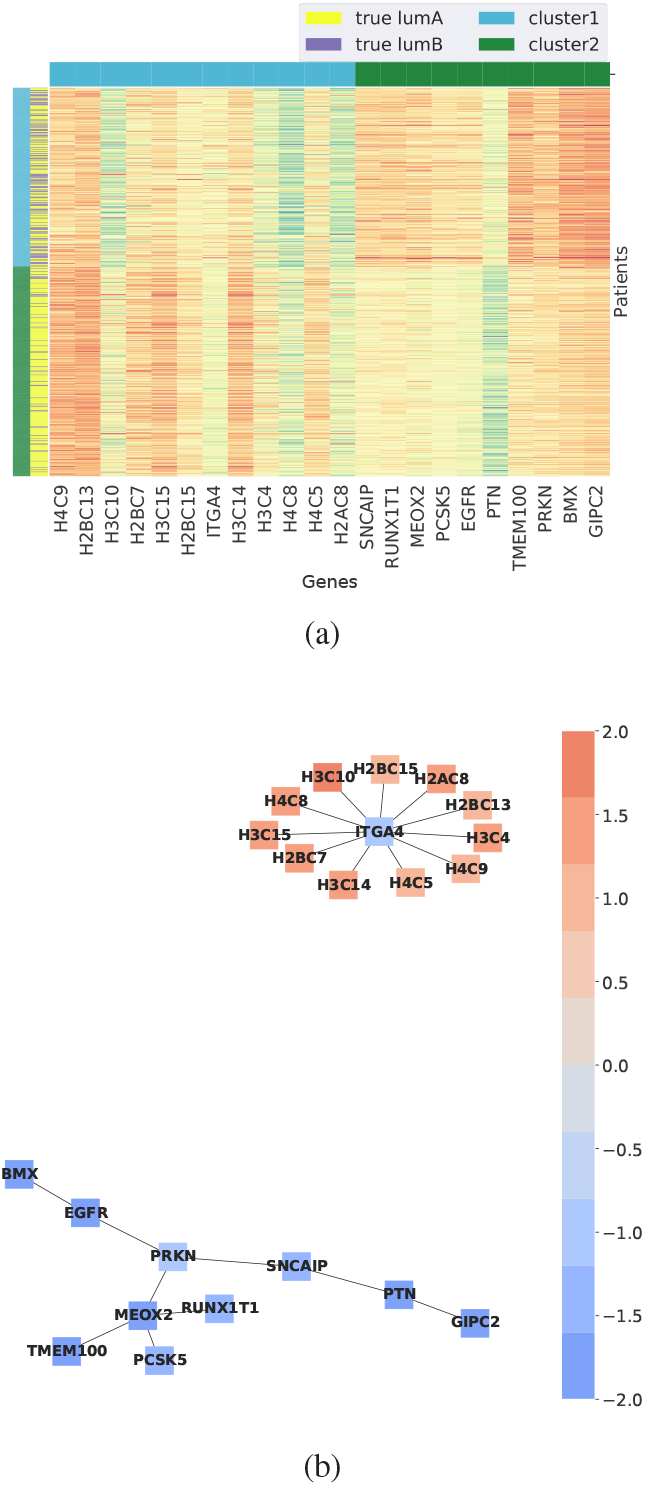
a) Biclusters for patients with luminal A and luminal B breast cancer subtypes. b) Resulting subnetwork: nodes are colored with respect to their mean expressions in one of the clusters and thus nodes colored orange have very high average expression in the cluster and the blue nodes have very low expression values.

#### 4.2.3 Basal patients stratification

Bertucci *et al.*, 2012 characterised basal, also known as triple negative, breast cancer as the most challenging breast cancer subtype with poor prognosis despite relatively high chemosensitivity. Currently, there is no targeted therapy and no routine diagnostic procedure specifically for this subtype. Although no clinically relevant subgroups of the basal subtype are known, BiCoN achieved a clear separation into two subgroups (Figure 8).

**Fig. 8:**
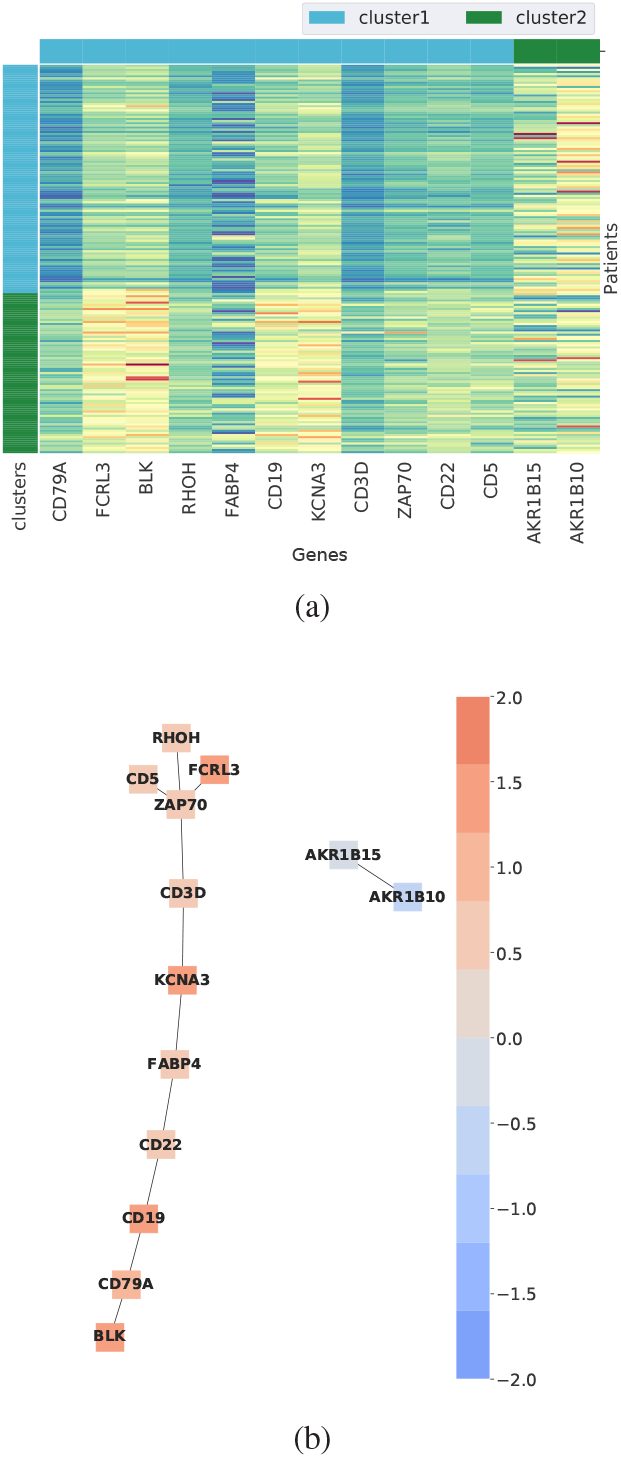
a) Biclusters for patients with basal breast cancer subtype.b) Resulting subnetwork: nodes are colored with respect to their mean expressions in one of the clusters and thus nodes colored orange have very high average expression in the cluster and the blue nodes have very low expression values.

Derived subnetworks show robust correlation with immune system response functions which is reasonable given that tumour samples are infiltrated with leukocytes. All 3 enriched pathways (Primary immunodeficiency, Hematopoietic cell lineage, B cell receptor signaling pathway) have a direct connection to the immune response (Figure S2(a) Supplementary Material). Molecular function enrichment also confirms the relation between the selected genes and immune response (Figure S2(b) Supplementary Material). Cell type deconvolution analysis with xCell shows a high correlation of the clusters with aDC, CD4+ memory T-cells, B-cells, CD8+ T-cells and other immune response related cells (Supplementary material Figures S4(b) and S5(b)). Similar to the results in luminal patients, our results indicate that basal breast cancer patients can be clustered by the contribution of tumor-infiltrating leukocytes, which is a key factor for prognosis and treatment via immunotherapy.

## 5 Conclusion and Outlook

Classical biclustering methods were shown to perform sub-optimally when non-intersecting, large patient subgroups are of interest as is often the case in patient stratification. Clustering methods, on the other hand, are more suited for this task, but they use the whole gene set and do not provide a mechanistic explanation of patient stratification. Therefore BiCoN is uniquely suited to cluster patients along with extracting fixed-size subnetworks capable of mechanistically explaining the patient stratification. Moreover, simultaneous clustering of gene expression and networks makes BiCoN robust to noise and more robust to batch effect than typical clustering and biclustering methods.

BiCoN leverages molecular interaction networks in the analysis of gene expression data to faithfully produce known subtypes as well as novel, clinically relevant patient subgroups, as we could demonstrate using data from TCGA. We stress that BiCoN and the concept of network-constrained biclustering are not limited to gene expression data or protein-protein interaction networks. We plan to apply BiCoN to other types of omics data such as DNA methylation, copy number variation or single nucleotide polymorphisms. We envision BiCoN to be useful for single-cell RNA-seq data for uncovering differences in signalling between clusters of cells and for the discovery of novel cell types. BiCoN, which is available as a web-interface and a PyPI package, has great potential to enhance our understanding of diseases, cellular heterogeneity and putative drug targets.

## Acknowledgements

The results shown here are in whole or part based upon data generated by the TCGA Research Network: https://www.cancer.gov/tcga.

We thank Quirin Heiß for his contributions to the source code of the web-interface.

## References

Aarts, E. et al. (2003). Local search in combinatorial optimization. Princeton University Press.

Alcaraz, N. et al. (2012). Efficient key pathway mining: combining networks and omics data. Integrative Biology, 4(7), 756–764.

Alcaraz, N. et al. (2017). De novo pathway-based biomarker identification. Nucleic Acids Res., 45(16), e151.

Aran, D. et al. (2017). xCell: digitally portraying the tissue cellular heterogeneity landscape. Genome Biol., 18(1), 220.

Banzhaf, W. et al. (1998). Genetic programming: an introduction, volume 1. Morgan Kaufmann San Francisco.

Barabási, A.-L. and Albert, R. (1999). Emergence of scaling in random networks. science, 286(5439), 509–512.

Batra, R. et al. (2017). On the performance of de novo pathway enrichment. NPJ systems biology and applications, 3(1), 6.

Berg, J. et al. (2002). Defects in signaling pathways can lead to cancer and other diseases. Biochemistry. 5th Edition. New York: WH Freeman, Section, 15.

Bertucci, F. et al. (2012). Basal breast cancer: a complex and deadly molecular subtype. Curr. Mol. Med., 12(1), 96–110.

Chen, C. et al. (2011). Removing batch effects in analysis of expression microarray data: an evaluation of six batch adjustment methods. PloS one, 6(2).

Cheng, Y. and Church, G. M. (2000). Biclustering of expression data. Proc Int Conf Intell Syst Mol Biol, 8, 93–103.

Eberhart, R. and Kennedy, J. (1995). Particle swarm optimization. In Proceedings of the IEEE international conference on neural networks, volume 4, pages 1942–1948. Citeseer.

Gillis, J. et al. (2014). Bias tradeoffs in the creation and analysis of protein– protein interaction networks. Journal of proteomics, 100, 44–54.

Goh, W. W. B. et al. (2017). Why Batch Effects Matter in Omics Data, and How to Avoid Them. Trends Biotechnol., 35(6), 498–507.

Hartigan, J. A. (1972). Direct Clustering of a Data Matrix. Journal of the American Statistical Association, 67(337), 123–129.

Lazar, C. et al. (2012). Batch effect removal methods for microarray gene expression data integration: a survey. Briefings in bioinformatics, 14(4), 469–490.

Lazzeroni, L. and Owen, A. (2002). Plaid models for gene expression data. Statistica sinica, pages 61–86.

Li, G. et al. (2009). Qubic: a qualitative biclustering algorithm for analyses of gene expression data. Nucleic acids research, 37(15), e101–e101.

Liu, Y. et al. (2014). A network-assisted co-clustering algorithm to discover cancer subtypes based on gene expression. BMC bioinformatics, 15(1), 37.

Luo, J. et al. (2010). A comparison of batch effect removal methods for enhancement of prediction performance using maqc-ii microarray gene expression data. The pharmacogenomics journal, 10(4), 278–291.

Matthews, B. W. (1975). Comparison of the predicted and observed secondary structure of t4 phage lysozyme. Biochimica et Biophysica Acta (BBA)-Protein Structure, 405(2), 442–451.

Morrison, J. L. et al. (2005). GeneRank: using search engine technology for the analysis of microarray experiments. BMC Bioinformatics, 6, 233.

Nielsen, T. O. et al. (2010). A comparison of PAM50 intrinsic subtyping with immunohistochemistry and clinical prognostic factors in tamoxifen-treated estrogen receptor-positive breast cancer. Clin. Cancer Res., 16(21), 5222–5232.

Nygaard, V. et al. (2016). Methods that remove batch effects while retaining group differences may lead to exaggerated confidence in downstream analyses. Biostatistics, 17(1), 29–39.

Parker, J. S. et al. (2009). Supervised risk predictor of breast cancer based on intrinsic subtypes. J. Clin. Oncol., 27(8), 1160–1167.

Prelić, A. et al. (2006). A systematic comparison and evaluation of biclustering methods for gene expression data. Bioinformatics, 22(9), 1122–1129.

Prelic, B. S. & Zimmermann, P. (May 2006). A systematic comparison and evaluation of biclustering methods for gene expression data. Bioinformatics, 22, 1122–1129.

Rousseaux, S. et al. (2013). Ectopic activation of germline and placental genes identifies aggressive metastasis-prone lung cancers. Science translational medicine, 5(186), 186ra66–186ra66.

Shi, J. and Malik, J. (2000). Normalized cuts and image segmentation. IEEE Transactions on pattern analysis and machine intelligence, 22(8), 888–905.

Stützle, T. (2009). Ant colony optimization. In M. Ehrgott, C. M. Fonseca, X. Gandibleux, J.-K. Hao, and M. Sevaux, editors, Evolutionary Multi-Criterion Optimization. Springer Berlin Heidelberg.

Stutzle, T. and Hoos, H. (1997). Max-min ant system and local search for the traveling salesman problem. In Proceedings of 1997 IEEE international conference on evolutionary computation (ICEC’97), pages 309–314. IEEE.

Stützle, T. and Hoos, H. (1999). The max-min ant system and local search for combinatorial optimization problems. In Meta-heuristics, pages 313–329. Springer.

Tanay, A. et al. (2002). Discovering statistically significant biclusters in gene expression data. Bioinformatics, 18 Suppl 1, S136–144.

Thangavelu, S. et al. (2019). Feature selection in cancer genetics using hybrid soft computing. pages 734–739.

Voorhees, E. M. (1986). Implementing agglomerative hierarchic clustering algorithms for use in document retrieval. Information Processing & Management, 22(6), 465–476.

Yu, G. et al. (2017). Network-aided Bi-Clustering for discovering cancer subtypes. Sci Rep, 7(1), 1046.

