## Supplementary for "BiCoN: Network-constrained biclustering of patients and omics data"

### Supplementary material

#### Algorithm description

Each of the methods used in the pseudo-code (Algorithm 1) is briefly outlined below. We use the following notation while referring to genes and patients:

- $n$ - number of genes, all gene IDs are mapped to the range [0,n-1]
- $m$ - number of patients, all patient IDs are mapped to the range [n,n+m-1]
- w.l.o.g. matrices used for transition through the graph including the heuristic information matrix ($H$), the pheromone matrix ($T$), the transition probability matrix ($P$) and the cost matrix ($C$) have dimensionality $(n+m) \times (n+m)$. Their row and column indices correspond to gene and patient IDs

##### Functions

The following functions are used in the Algorithm 1.

**HeuristicMat**: The transition probability depends on a pheromone level and an initially given heuristic information (Equation 2). The symmetric matrix $H^{(n+m),(n+m)}$ consists of all pairwise similarities between all nodes of the graph $J$ (both patients and genes). The pairwise similarity between two nodes is defined as follows depending whether the nodes belong to a set of patients (the $P$ set) or a set of genes (the $V$ set):

$$H[i,j] = \begin{cases} X[i,j] & \text{if } i \in V, j \in P \\ cor(X[,i], X[,j]) & \text{if } i \in P, j \in P \\ cor(X[i,], X[j,]) & \text{if } i \in V, j \in V \text{\& the genes are adjacent} \\ 0 & \text{if } i \in V, j \in V \text{\& the genes are NOT adjacent} \\ 0 & \text{if } i = j \end{cases} \quad (1)$$

All similarity values are scaled to the range [1,10] and it is ensured that they are on the scale as the pheromone values. The main diagonal is set to 0 in order to avoid stagnation on a single node.

**CostCalc**: Ants are restricted in the number of steps they can take. We allow ants to travel longer if they choose highly similar nodes. To achieve this, we define a cost matrix $C = 1 - H \times \frac{1}{10}$ which specifies the cost for a transition between two nodes. In total, each ant can spend a fixed value $cl$ (see Algorithm 1). We tested various values for $cl$ and found that its choice has little influence. Thus, we set $cl$ automatically depending on the desired solution size.

**InitialPher**: The initial pheromone level $T$ is determined by (as suggested in for Min-Max ACO): $T = \mathbf{1} \cdot \mathbf{1}^T \cdot score_{max}$

where $\mathbf{1} \cdot \mathbf{1}^T$ stands for the matrix of all ones in $\mathbb{R}^{(n+m),(n+m)}$.

**ProbUpd**: We use the classic ACO probability update approach where the transition probability represents a scaled product of the pheromone matrix $T$ and the heuristic information $H$. Both matrices have corresponding importance levels $\alpha$ (pheromone significance) and $\beta$ (heuristic information significance) which allow balancing the influence of $T$ and $H$. Our experiments show that the best performance is achieved when $\alpha = \beta = 1$ as by design $H$ and $T$ matrices are scaled to the same range.

The probability of transitioning from node $i$ to node $j$ for a patient $s$ is thus defined as:

$$P[i,j]_s = \begin{cases} \frac{T[i,j]^\alpha H[i,j]^\beta}{\sum_{j \in N_s} T[i,j]^\alpha H[i,j]^\beta} & \text{if } j \in N_s \\ 0 & \text{otherwise} \end{cases} \quad (2)$$

Where $N_s$ - is the domain of an ant. $N_s$ is computed for each patient with a function *SearchRad* (see bellow).

**SearchRad**: Determines feasible transitions for a given ant. Each ant can perform the search in the domain linked to a patient. We achieve this during the

*RandomWalk* procedure by assigning an ant to every patient, i.e. we limit the search space of an ant by allowing transitions only to nodes which are similar to the starting node. As a similarity measure we use the heuristic information matrix:

$$N_s = [i|(H[s,i] > \bar{H}[s,]], \text{ for } s = n, n+1, \ldots, n+m, \text{ for } i = 0, 1, \ldots, n+m \quad (3)$$

**RandomWalk**: Performed for each patient according to Equation (2). An ant is travelling in the domain defined by $N$ until the cost limit is reached.

**ClusterPatients**: We cluster patients with a k-means algorithm considering only genes visited during the random walks.

**ClusterGenes**: Given patient clusters and genes that were chosen for each patient by the ants, we split genes into groups depending on the patient for which they were picked. An example of this procedure is given bellow.

In Table S1 we listed patients and genes selected during the random walk as well as a cluster of each patient determined with *ClusterPatients* function.

At the next step, we can split genes with respect to clusters where they have appeared (Table S2). Naturally, some genes have appeared in more than one cluster which means that even though they are relevant for both groups of patients, they are not relevant for the clustering (as Gene 1 in the Table S2) and therefore we delete them from both clusters.

| **Patient** | **Corresponding random walk** | **Cluster** |
|---|---|---|
| Patient1 | gene1,gene2 | 1 |
| Patient2 | gene2,gene3 | 1 |
| Patient3 | gene1,gene4 | 2 |
| Patient4 | gene4, gene5 | 2 |

Table S1: Input for the genes clustering procedure: each patient has a linked random walk results and a cluster determined at the previous step of the algorithm

| **Gene** | **Cluster** |
| --- | --- |
| ~~Gene1~~ | ~~1,2~~ |
| Gene2 | 1 |
| Gene3 | 1 |
| Gene4 | 2 |
| Gene5 | 2 |

Table S2: Output of the genes clustering procedure: each gene that was in at least one random walk is assigned to at least one cluster. Then, genes that appeared to be in more than one cluster get filtered out

**NetReduce**: After ACO is complete, the selected genes form two (in the current implementation, $c$ in the general case) subnetworks. Those subnetworks might be larger than the user-provided maximum subnetwork size $L_{max}$ and hence we check, if we can decrease the subnetwork size by removing uninformative genes. We reduce the size of the resulting subnetworks by iteratively removing singletons or nodes with low node degree and smallest average expression difference between patient clusters. The procedure is repeated until we obtain $c$ networks of at least $L_{min}$ genes and at most $L_{max}$ genes.

**Score**: Each solution is rated by the value of the objective function (Equation 1 in the main paper). With respect to gene expression, our overall goal is to obtain $c$ subnetworks with high average expression levels inside their patients group (inner expression) and a low average expression level outside of it (outer expression). To achieve this, we chose an objective function that measures the difference between inner expression and outer expression and then scales it with respect to how well the networks are connected inside their cluster. Thus, we can ensure that disconnected nodes with even very high difference in expression will not be considered by the algorithm at all since their score will be equal to 0.

**PherUpd**: After the best solution is determined, we update the pheromone matrix $T$ based on the optimal score $s_{max}$:

$$T[i,j] = \rho * T[i,j] + \Delta T[i,j] \tag{4}$$

where $\rho$ is an evaporation rate and $\Delta T[i,j]$ is calculated as:

$$\Delta T[i,j] = \begin{cases} s_{max} & \text{if } i,j \text{ belong to the same cluster} \\ 0 & \text{otherwise} \end{cases} \tag{5}$$

After the update, we enforce $T[i,j] \geq 0 \forall i,j$.

As a termination criterion, we consider the difference between the best and the average score of each run. Thus, if $s_{max} - \bar{s} < \varepsilon$, the algorithm terminates and the best solution is reported.

**LocalSearch**: A user can additionally run local search optimization on top of a fully converged ACO solution. Local search ensures that a solution is locally optimal by making small changes to the retrieved subnetworks (node insertion, deletion or substitution) and evaluating contribution to the objective function. When local improvements are not possible anymore, the algorithm returns an improved solution.

#### Supplementary visualization material

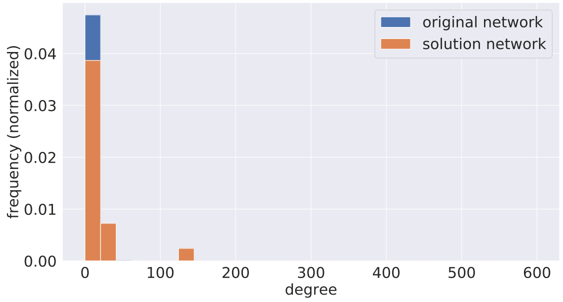


Figure S1: Solution subnetworks degree distribution after 10 runs against the input network degree distribution.

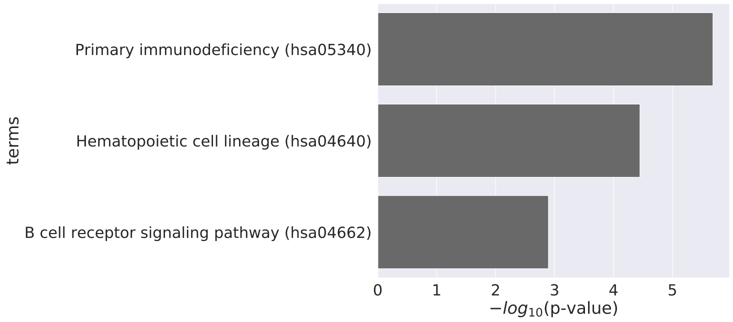


(a)

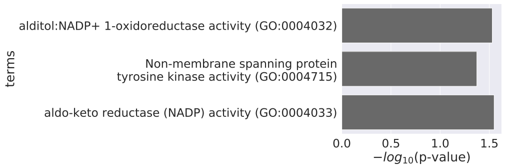


(b)

Figure S2: **a)** KEGG gene enrichment for patients with basal breast cancer subtype (based on the solution genes). **b)** Molecular Function GO term enrichment based on the solution genes.

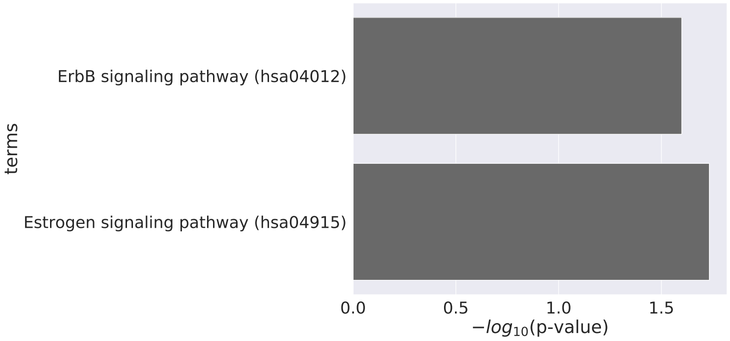


Figure S3: Enrichment of the solution genes for patients with basal and luminal breast cancer.

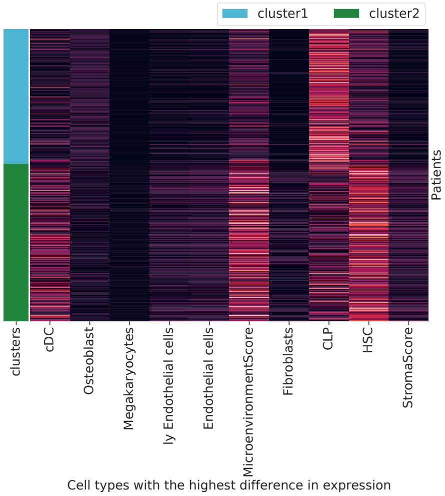


(a)

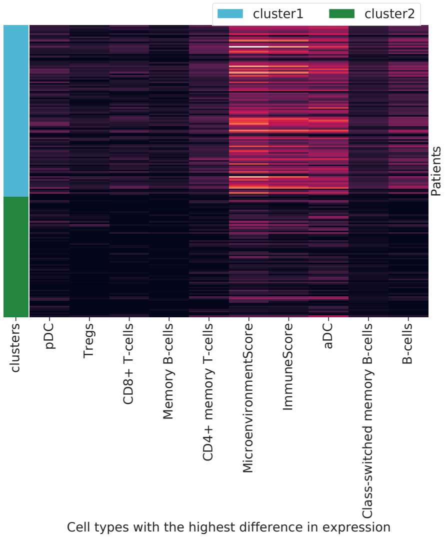


(b)

Figure S4: Top 10 cells with the highest difference between achieved clusters for patients with luminal cancer subtype **(a)** and with basal cancer subtype **(b)**.

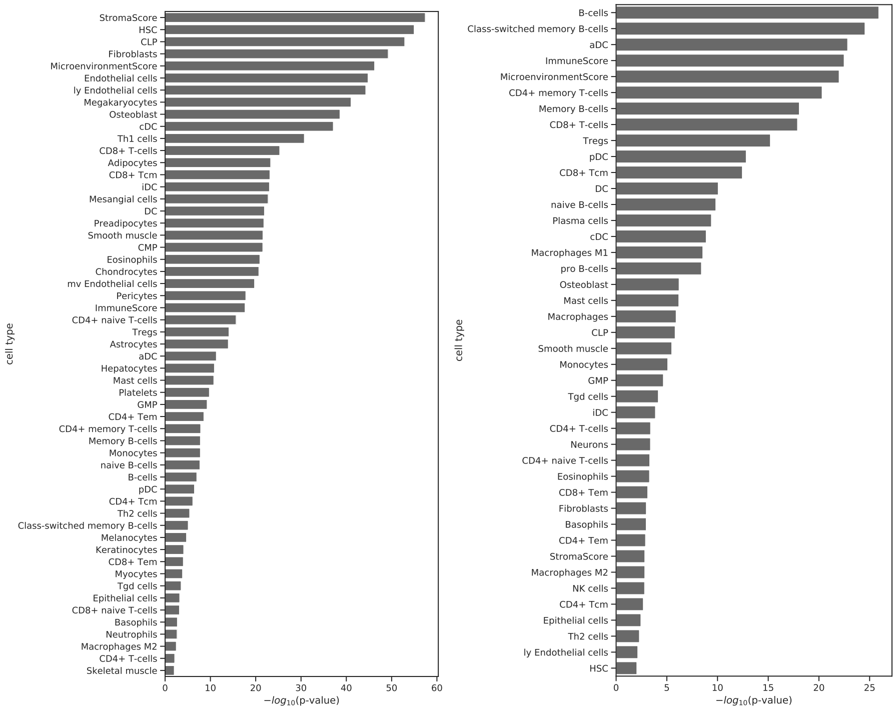


(a) (b)

Figure S5: T-test -log10 p-values for scores comparison between two achieved clusters for patients with luminal breast cancer subtype **(a)** and for patients with basal breast cancer subtype **(b)**.
